# Antibiotic production in *Streptomyces* is organized by a division of labour through terminal genomic differentiation

**DOI:** 10.1101/560136

**Authors:** Zheren Zhang, Chao Du, Frederique de Barsy, Michael Liem, Apostolos Liakopoulos, Gilles P. van Wezel, Young H. Choi, Dennis Claessen, Daniel E. Rozen

**Affiliations:** Institute of Biology, Leiden University, Sylviusweg 72, 2333 BE, Leiden, The Netherlands

## Abstract

One of the hallmark behaviors of social groups is division of labour, where different group members become specialized to carry out complementary tasks. By dividing labour, cooperative groups of individuals increase their efficiency, thereby raising group fitness even if these specialized behaviors reduce the fitness of individual group members. Here we provide evidence that antibiotic production in colonies of the multicellular bacterium *Streptomyces coelicolor* is coordinated by a division of labour. We show that *S. coelicolor* colonies are genetically heterogeneous due to massive amplifications and deletions to the chromosome. Cells with gross chromosomal changes produce an increased diversity of secondary metabolites and secrete significantly more antibiotics; however, these changes come at the cost of dramatically reduced individual fitness, providing direct evidence for a trade-off between secondary metabolite production and fitness. Finally, we show that colonies containing mixtures of mutant strains and their parents produce significantly more antibiotics, while colony-wide spore production remains unchanged. Our work demonstrates that by generating mutants that are specialized to hyper-produce antibiotics, streptomycetes reduce the colony-wide fitness costs of secreted secondary metabolites while maximizing the yield and diversity of these products.

## Introduction

Social insects provide some of the most compelling examples of divisions of labour, with extremes in morphological differentiation associated with highly specialized functions and reproductive sterility in all colony members except the queen^1^. However, conditions that select for division of labour are not limited to animals and it has become increasingly clear that microbes offer unique opportunities to identify and study the mechanistic underpinnings of divisions of labour^2–8^. First, microbes are typically clonal, which helps ensure that a division of labour is favoured by kin selection^4^. Second, microbial populations are highly social, often cooperating to carry out coordinated behaviors like migration or biofilm formation that require the secretion of metabolically expensive public goods that can be shared among clonemates^9, 10^. If these conditions are met, and investment in public good secretion trades-off with fitness, divisions of labour are predicted to evolve^4, 11^.

Here we describe the cause and evolutionary benefits of a unique division of labour that has evolved in colonies of the filamentous actinomycete *Streptomyces coelicolor*. After germinating from unichromosomal spores, these bacteria establish multicellular networks of vegetative hyphae, reminiscent of fungal colonies^12–14^. Vegetative hyphae secrete a broad variety of public goods, such as chitinases and cellulases that are used to acquire resources, as well as a chemically diverse suite of antibiotics that are used to kill or inhibit competing organisms^15–17^. Streptomycetes are prolific producers of antibiotics, and are responsible for producing more than 50% of our clinically relevant antibiotics^18^. Although the terminal differentiation of *Streptomyces* colonies into vegetative hyphae (soma) and viable spores (germ) is well understood^19–21^, no other divisions of labour in these multicellular bacteria are known. However, opportunities for phenotypic differentiation are possible, because even though colonies begin clonally, they can become genetically heterogeneous because of unexplained high-frequency rearrangements and deletions in their large, ∼9 Mb linear chromosome ^22–25^. The work we describe shows that these two topics are intertwined. Briefly, we find that genomic instability causes irreversible genetic differentiation within a subpopulation of growing cells. This in turn gives rise to a division of labour that increases the productivity and diversity of secreted antibiotics and increases colony-wide fitness.

## Results

### Genome instability and phenotypic heterogeneity are coupled

Genome instability and phenotypic heterogeneity have been observed in several *Streptomyces* species^26–32^, but there are no explanations for the evolution or functional consequences of this extreme mutability. To begin addressing this question, we quantified the phenotypic heterogeneity arising within 81 random single colonies of *S. coelicolor* M145 by harvesting the spores of each of these colonies and then replating the collected spores onto a new agar surface. Although most progeny are morphologically homogeneous and similar to the wild-type, strikingly aberrant colonies (Figure 1a) arise at high frequencies (0.79 +/-0.06%, mean +/-SEM, ranging from 0% to 2.15%, n = 81) (Figure 1a). Similarly high rates were obtained on two minimal media (MM: 2.13 +/-0.14% and MM+CA: 5.13 +/-0.37%, mean +/-SEM, n = 30 and n = 40 respectively) (Figure S1) thereby ruling out the possibility that these mutations are an artifact of rapid growth on rich resources. To determine the heritability of these aberrant phenotypes, we restreaked 15 random colonies from different plates onto a new agar plate which revealed remarkably variability in colony morphology (Figure 1b). Rather than reverting to the wild-type (WT) morphology, as would be anticipated if the initial heterogeneity were due to phenotypic plasticity or another form of bistability, the colonies derived from mutant colonies are themselves hypervariable, giving rise to up to nine diverse phenotypes from any single colony. Thus, in the course of two cycles of colony outgrowth, an array of colony types emerged that differ in size, shape and colour (Figure 1b). Because our ability to discern colony heterogeneity is limited to only a few visually-distinct phenotypic characters, we assume that these estimates of diversity are lower than their true level of occurrence.

**Figure 1:**
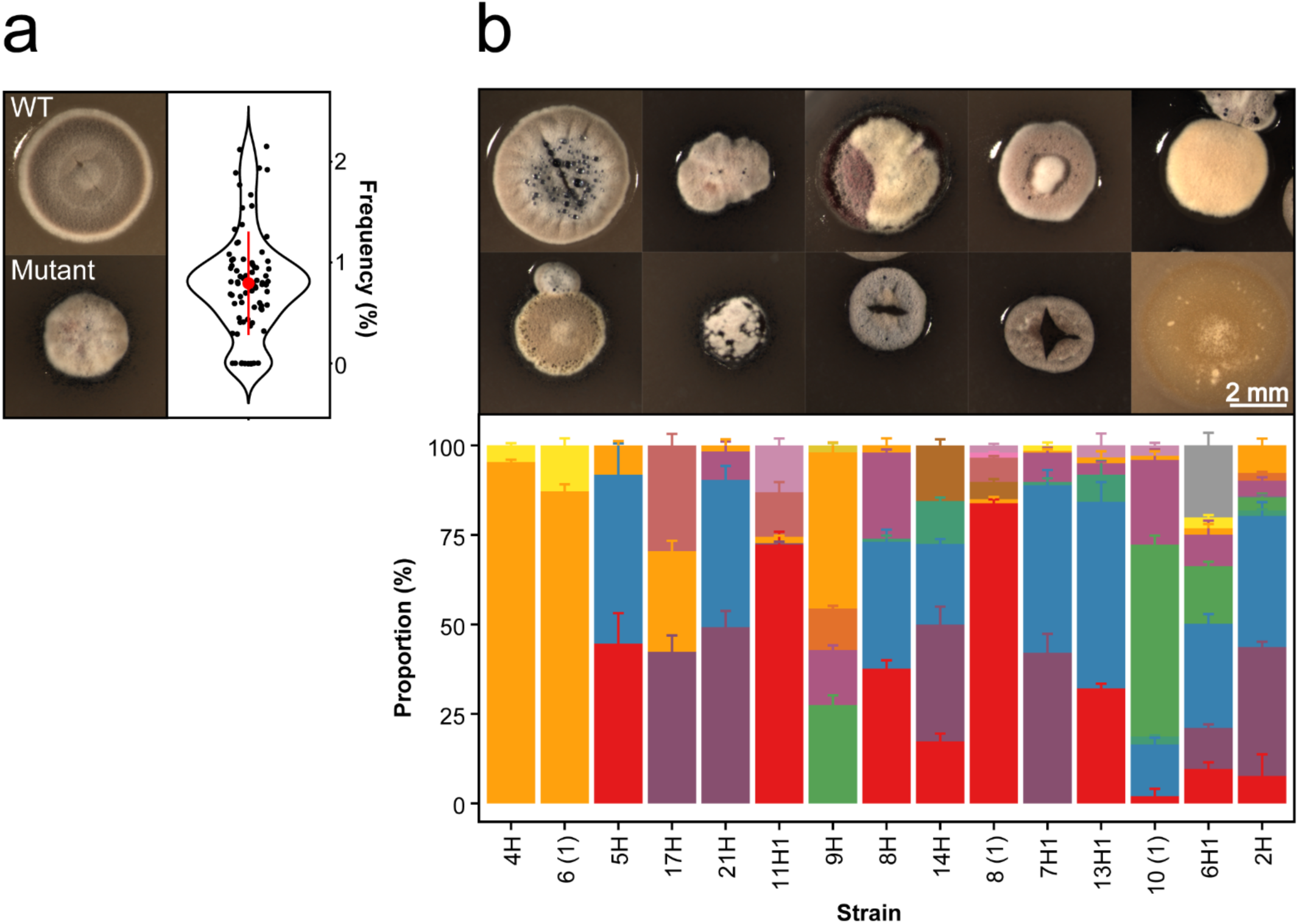
The emergence of phenotypic heterogeneity in colonies of *Streptomyces coelicolor*. (a) Wild-type (top) and mutant (bottom) colonies, and the frequency that mutants emerge from WT colonies (right). (B) Phenotypically diverse progeny (top) emerge after re-streaking mutant colonies that vary in size, shape and pigmentation. Representative colonies are shown. The bottom graph depicts the range of distinct morphologies that emerge after re-streaking 15 random colonies. Each color represents a distinct colony phenotype.

Using whole-genome sequencing of 8 random mutants we confirmed that these isolates contained profound chromosomal changes. As shown in Figure 2a, large genome deletions were observed at the chromosome ends in all 8 strains. In 3 cases we found an ∼297 kb amplification on the left chromosomal arm flanked by the Insertion Sequence IS1649, encoced by SCO0091 and SCO0368. Average sequence coverage of the amplified region suggests it contains between 2 and 15 copies of this amplification (Figure 2a and S2). Sequencing results were expanded using pulsed-field gel electrophoresis (PFGE) analysis of 30 mutant isolates (Figure 2b and S3). Consistent with our sequencing results, this analysis revealed that mutants contained variably sized deletions of up to ∼ 240 kb or ∼ 872 kb on the left chromosome arm and up to 1.6 Mb on the right chromosome arm, deleting more than 1,000 genes. Additionally, 8/30 strains contained the same large amplification between copies of IS1649 as noted above. Interestingly, these strains are conspicuously yellow, which might be caused by the overproduction of carotenoids due to the amplification of the *crt* gene cluster (SCO0185-0191;^33–35^). In addition to this and other phenotypic effects associated with these changes that are discussed below, deletions to the right chromosome arm cause the loss of two loci, *argG* (SCO7036) and *cmlR1*(SCO7526)*/cmlR2*(SCO7662), that result in two easily scorable phenotypes: arginine auxotrophy and chloramphenicol susceptibility, respectively. Scoring these phenotypes allows rapid determination of the minimal size of the deletion on the right chromosome arm in the absence of molecular characterization. Chloramphenicol susceptibility indicates a deletion of at least 322 kb, while the addition of arginine auxotrophy indicates a deletion of at least 843 kb (Figure 2b).

**Figure 2:**
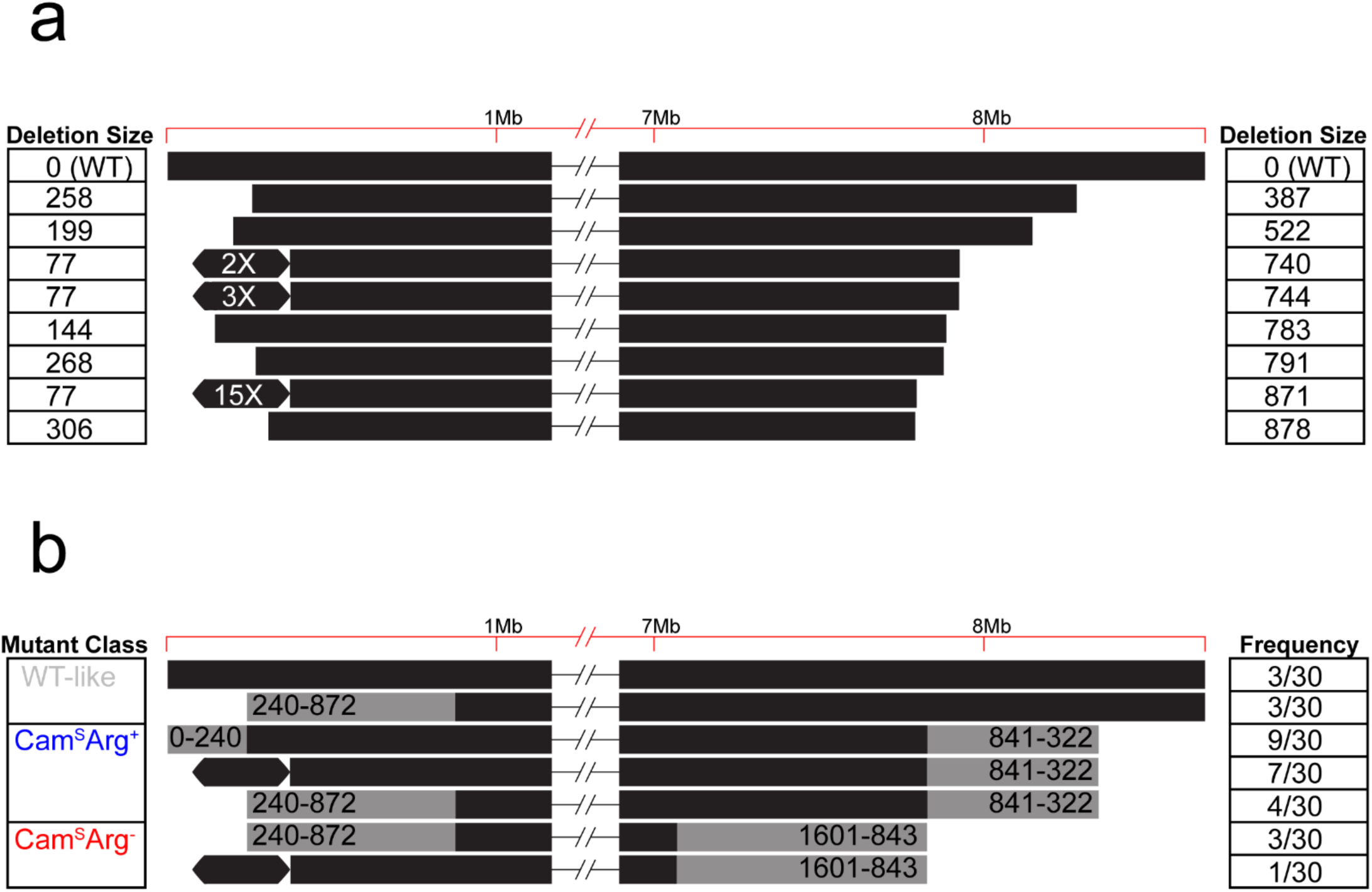
Genome diversity of mutant colonies determined from (a) whole-genome sequencing and (b) Pulsed-field gel electrophoresis (PFGE). Values in (a) correspond to the size (in kb) of genome deletions while the hexagons represent an ∼297 kb genome amplification. Each line in (b) depicts the range of deletion sizes (grey) in each mutant class, together with their respective frequencies from 30 sampled mutant strains.

### Mutants increase the production and diversity of antibiotics

Mutant strains were conspicuously pigmented when compared to their parental wild-type strains (Figure 1). Because several antibiotics produced by *S. coelicolor* are pigmented, namely actinorhodin, prodigines and coelimycin P which are blue, red and yellow, respectively, we tested if mutant strains had altered secondary metabolite and inhbitory profiles. Secreted metabolites from mutant and WT strains grown on agar surfaces were analyzed using quantitative ^1^H NMR profiling^36, 37^. Principal component analysis (PCA) (Figure 3a) supports the partition of strains into three well-separated groups: wild-type and wild-type-like strains, and then two clusters of mutant isolates. In each case, groupings corresponded to the size of genomic lesions mentioned above. More specifically, strains grouping in the wild-type and wild-type-like cluster are chloramphenicol resistant (Cam^R^) and arginine prototrophic (Arg^+^), while those clustering within the blue ellipse were chloramphenicol susceptible (Cam^S^) and protrophic for arginine (Arg^+^), and those in the red-ellipsed cluster were susceptible to chloramphenicol (Cam^S^) and auxotrophic for arginine (Arg^−^). To assess if genomic deletions affected antibiotic biosynthesis, we used mass spectrometry-based quantitative proteomics on five representative strains from the two mutant clusters. This analysis revealed that the biosynthetic pathways for actinorhodin, coelimycin P1 and calcium-dependent antibiotic (CDA) were significantly upregulated in all mutants (Figure 3b, c, S4 and Table S1). Because the expression level of biosynthetic enzymes directly correlates with antibiotic production^38^, these MS results are consistent with increased antibiotic production in these strains (Figure 3b, c, d and S4). In addition to antibiotic biosynthesis clusters, pathways regulating arginine and pyrimidine biosynthesis were also increased in both arginine auxotrophic strains (Figure S4b and Table S1)^39^. No antibiotic-related proteins were down-regulated in this analysis.

**Figure 3:**
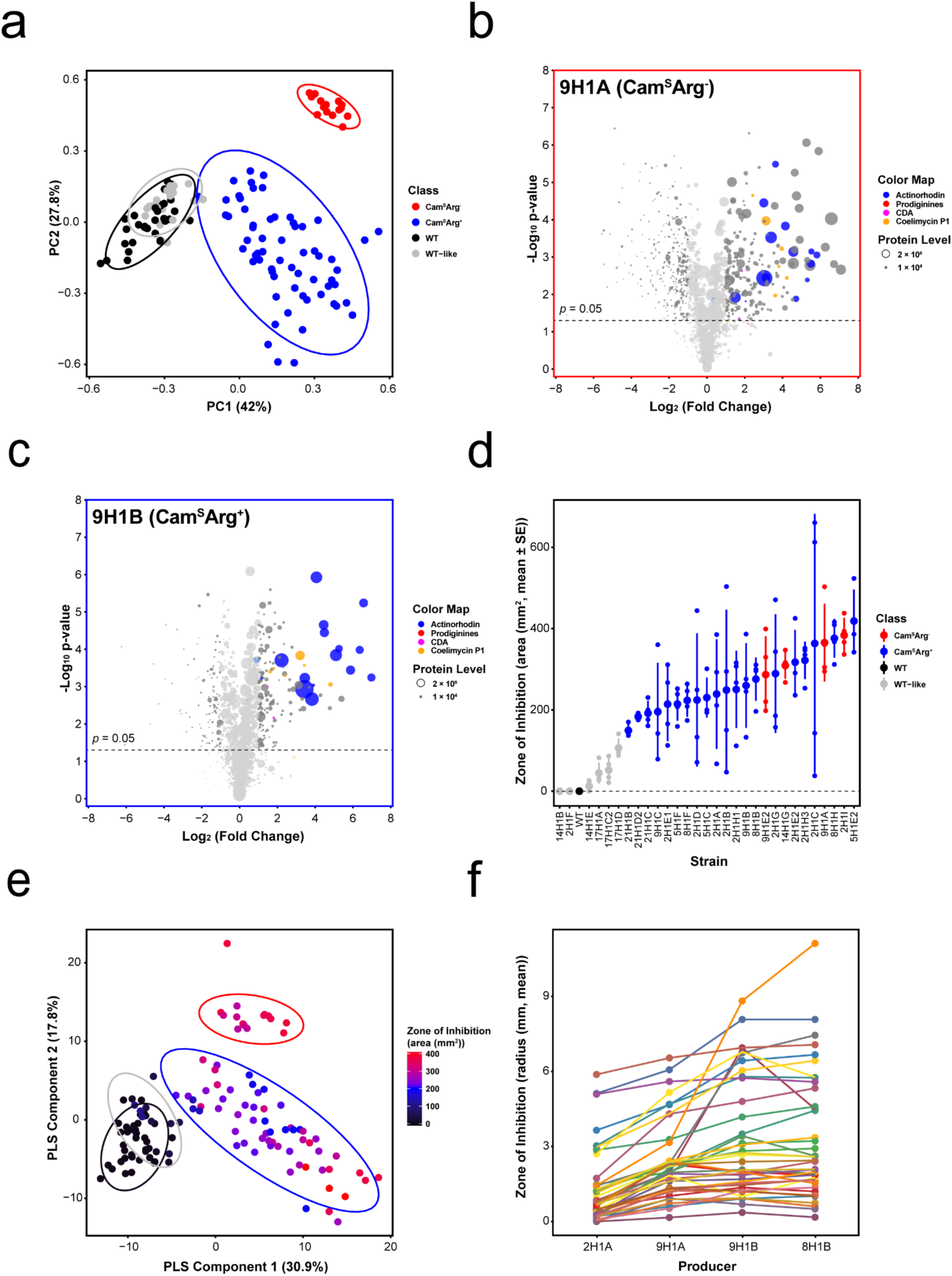
Secondary metabolite production in mutant strains determined by ^1^H NMR (a and e), quantitative proteomics (b and c) or zones of inhibition on *B. subtilis* (d) or 40 different natural streptomycete isolates (f). (a) PCA plot of ^1^H NMR data. Each cluster enclosed in a colored ellipse (with 95% CI) corresponds to a mutant class with a different phenotype and degree of genomic instability: WT-like strains (in grey), Cam^S^Arg^+^ strains (in blue) and Cam^S^Arg^−^ strains (in red). (b, c) Volcano plots of MS-based quantitative proteomics of two representative strains 9H1A (Cam^S^Arg^−^) (b) and 9H1B (Cam^S^Arg^+^) (c). Protein level is indicated by the size of the dot and genes with ≦ 2-fold change and/or *p* > 0.05 are greyed out. (d) Zones of inhibition of each strain when grown with a *B. subtilis* soft-agar overlay. Colors represent the same mutant classes as in (a). The large dot represents the mean of four replicates while error bars represent the standard error. (e) PLS plot of ^1^H NMR data partitioned by the same clusters as in (a). The heat map indicates the size of the zone of inhibition on *B. subtilis.* (f) Zones of inhibition of 4 representative mutant strains with an overlay of 40 different natural streptomycetes, each represented by a different line. Statistics are given in the main text.

We next asked if these different metabolic and proteomic profiles translated to differences in biological activity, specifically the ability to inhibit the growth of other bacteria. Thirty mutant strains were grown on agar plates and then covered with a soft-agar overlay containing *B. subtilis*. Inhibition was visualized as an absence of growth surrounding the mutant colony, and the extent of inhibition was determined from the size of the inhibition zone. As shown in Figure 3d, all but three WT-like mutant strains produced significantly larger zones of inhibition than the WT strain (One-tailed t-tests, all *p* < 0.05). In addition, we observed significant heterogeneity among mutant strains in halo size (One-Way ANOVA, F_29,90_ = 5.45, *p* < 0.001).

To test if the increased inhibition we observed against *B. subtilis* was correlated with the ^1^H NMR profiles, we used a partial least squares (PLS) regression (Figure 3e)^37^. This showed that the separation into different groups significantly correlates with halo size (Q^2^ = 0.879) which was further validated by both permutation-tests and ANOVA of cross-validated residuals (CV-ANOVA, F_8,116_ = 104.443, *p* < 0.001). To identify possible compounds that are overproduced in mutants compared to WT, we identified several ^1^H NMR signals that varied across strains and strongly correlated with the size of the zone of inhibition against *B. subtilis* (Table S2); notable among these are several aromatic signals which correspond to actinorhodin, consistent with our proteomic analyses (Figure 3b and c).

Phenotypic results indicate that mutant strains produce more antibiotics than their wild-type parent when assayed against a single bacterial target, as anticipated given our NMR and proteomic results. However, they do not distinguish if strains can be further partitioned on the basis of which other species they inhibit. Score plots of principal component analysis based on ^1^H NMR signals reveal clear separation between mutant clones within and between clusters (Figure 3a), suggesting that their inhibitory spectra may vary. In addition, quantitative proteomic data show that different strains vary in their production of known antimicrobials. To test this, we measured the ability of four mutant clones to inhibit 48 recently isolated *Streptomyces* strains^40^. *Streptomyces* targets were chosen because these are likely to represent important competitors for other streptomycetes in soil environments. At least one of the four mutant strains produced a significantly larger halo than the wild-type strain against 40 out of 48 targets, indicating increased inhibition (Figure 3f). More interestingly, for these 40 targets, we observed significant differences between the mutant strains themselves. We found differences in the size of the zone of inhibition on different target species (Two-Way ANOVA, F_39,117_ = 21.79, *p* < 0.001) as well as a significant interaction between mutant strain and the target species (Two-Way ANOVA, F_117,320_ = 5.75, *p* < 0.001), indicating that the inhibitory profile of each mutant strain is distinct from the others. Together these results reveal that mutants arising within colonies are not only more effective at inhibiting other strains, but that they are also diversified in who they can inhibit because their inhibition spectra do not overlap. They also suggest that the beneficiary of diversified antibiotic secretion is the parent strain, because competing bacteria are unlikely to be resistant to this broadened combination of secreted antimicrobials.

### Antibiotic production is coordinated by a division of labour

Having shown that *Streptomyces* colonies differentiate into distinct subpopulations that vary in their antibiotic production, we next asked how this differentiation affects colony fitness. To answer this, we measured the fitness of each mutant strain by quantifying the number of spores they produce when grown in isolation. As shown in Figure 4a, mutants produce significantly fewer spores than the wild-type strain (all *p* < 0.001) and, in extreme cases, as much as 10,000-fold less, with significant heterogeneity among strains (One-Way ANOVA: F_29,59_ = 132.57, *p* < 0.001). Importantly, the reduction in spore production is significantly negatively correlated with antibiotic production (F_1,29_ = 26.51, r^2^ = 0.478, *p* < 0.001) (Figure S5a). This provides evidence that antibiotic production is costly to *S. coelicolor* and that there is a direct trade-off between antibiotic production and reproductive capacity, possibly because energy is redirected from development to antibiotic production^41^. In addition, we observed a significant negative correlation between the size of the genome deletion and CFU (F_1,7_ = 12.32, r^2^ = 0.638, *p* = 0.0099) and a positive correlation between deletion size and bioactivity against *B. subtilis* (F_1,7_ = 37.97, r^2^ = 0.844, *p* < 0.001), suggesting that these phenotypes scale with the magnitude of genomic changes (Figure 4b).

**Figure 4:**
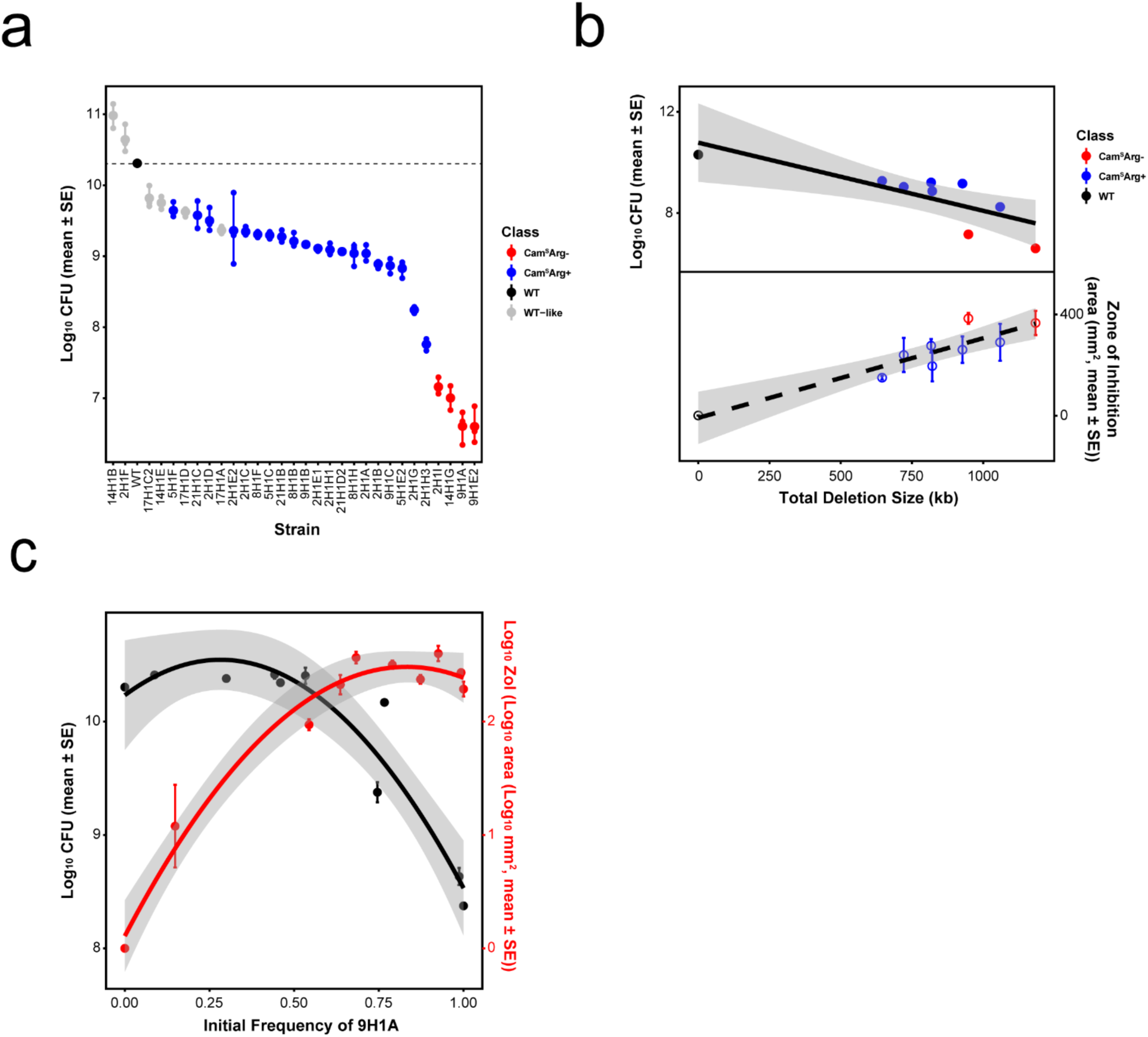
(a) Fitness (CFU) of mutant strains. (b) Decreases in genome size negatively correlate with fitness (top panel) and positively correlate with antibiotic production (lower panel). (c) Division of labour during co-culture of the WT and strain 9H1A at different starting frequencies. Increasing frequencies of 9H1A cause increased antibiotic production (F_2,7_ = 37.95, r^2^ = 0.916, *p* < 0.001) (red), but only negatively impact colony fitness at frequencies > ∼ 50% (F_2,13_ = 131.7, r^2^ = 0.953, *p* < 0.001) (black). Quadratic regression lines include the 95% CI.

To examine the effects of mutant strains on the colony as a whole, we mixed mutant strains with their parent at increasing frequencies and quantified colony-wide spore production and the ability of these mixtures to kill *B. subtilis*. Results of these experiments, shown in Figure 4c, support two important conclusions: 1) increasing fractions of mutants lead to increased antibiotic production; and 2) even though mutant strains have individually reduced fitness (Figure S5b), their presence within colonies has no effect on colony-wide spore production, until the mutant frequency exceeds > 50% of the total. We carried out the same assay with 3 additional mutant clones, but at fewer frequencies to estimate spore production, and observed concordant results (Figure S6): up to a frequency of ∼ 50%, mutant strains have no effect on colony-wide spore production, while each incremental increase in the frequency of these strains enhances colony-wide antibiotic output. These results indicate that the benefits of producing cells with chromosomal lesions are evident across a broad range of frequencies, but that even with extremely high mutation rates the costs to colony-wide fitness are minimal or entirely absent.

## Discussion

Streptomycetes are prolific producers of antibiotics, with genomes typically containing more than 20 secondary metabolite gene clusters that comprise more than 5% of their entire genome^34, 42, 43^. They invest heavily in these products and their biosynthesis and secretion is costly. Our results suggest that by limiting antibiotic production to a fraction of the colony through division of labour, *S. coelicolor* can eliminate the overall costs of biosynthesis while maximizing both the magnitude and diversity of their secreted antibiotics. Although this comes at a large individual cost, it increases group fitness by improving the ability for *S. coelicolor* to inhibit their competitors. Moreover, our results reveal that the range of conditions that select for a division of labour are quite broad, because colony-wide fitness is unaffected, even if mutant strains are as frequent as ∼ 50%.

Division of labour is predicted to be favoured in this system for several reasons. First, *Streptomyces* colonies emerge from a single spore and are clonal^19^. This, together with their filamentous mode-of-growth, ensures that costly individual traits can be maintained due to their indirect fitness benefits ^4, 5, 13^. And because resistance to the diversified antibiotic profile of mutant strains is unlikely to be present in competing strains, only the parent strain stands to benefit from their sacrifice. Second, the costs of antibiotic production via large and dedicated multi-step biosynthetic pathways, e.g. non-ribosomal peptide or polyketide synthases, are likely to be highest at the initiation of antibiotic production but diminish thereafter, meaning that producing cells become more efficient at making antibiotics through time^11^; furthermore, we show that antibiotic production trades-off with reproduction. Finally, many antibiotics are secreted, so the entire colony, but not susceptible competitors, can benefit from the protection they provide^44^.

Even if conditions predispose to a division of labour, there must still be a process that generates phenotypic heterogeneity. Our results show that in *Streptomyces*, this is caused by genomic instability that creates a subpopulation of cells within colonies that contain large deletions or amplifications at the termini^23, 25, 31^. Importantly, these mutations are severe and irreversible. Because strains, or portions of colonies, containing these deletions have significantly reduced fitness, they effectively behave like a sterile caste that provide direct benefits to the rest of the colony and receive little in return^1^. Indeed, even when the initial frequency of mutants in mixed colonies approaches 80%, their final frequency declines to less than 1% after one cycle of colony growth (Figure S7). This suggests that the division labour in *S. coelicolor* is re-established independently and differently in each colony, a mechanism that may help to maximize the diversity of secreted antibiotics.

It remains unclear if there are mechanisms regulating the size and frequency of chromosomal deletions and amplifications. One possibility is that these events are induced by external environmental conditions and that their rate is context dependent. Instability can be elevated by exposure to certain toxicants, e.g. mitomycin C or nitrous acid^45^, although no explicit stress was added in the experiments we report. It may also be increased during competition with other strains, a process that is known to alter the secretion of secondary metabolites^16, 46^. Another possibility is that deletions, and the benefits they bring for antibiotic production, are a fortuitous byproduct of the cell death that accompanies development^19^. By this argument, chromosome degradation would be regulated, but wouldn’t always be lethal. Although we have not confirmed this experimentally, it is likely that conserved amplifications result from the flanking copies of IS1649, which can facilitate intragenomic rearrangements^47^. In either case, the expectation is that increased antibiotic production results from the deregulation of biosynthetic clusters following the deletion of 100s of genes, many known to coordinate antibiotic biosynthesis^34^. And because deletions are stochastic, especially following the removal of protective telomeres at the ends of linear chromosomes, this would also cause antibiotic production to vary in different sections of the colony.

Our preliminary surveys have found similar levels of genomic instability in streptomycete strains we have freshly isolated from soil, suggesting that the division of labour we describe here is general. We are limited, however, in our ability to detect polymorphisms; color changes are conspicuous and are invariably associated with changes to pigmented secondary metabolites, but other secreted public goods may also become modified in these multicellular bacteria. Understanding which, if any other, public goods vary in the ways shown here is crucial because it will help to identify conditions that lead to a genetically encoded division of labour as compared to other forms of regulation that allow complex multicellular microbial systems to coordinate their behaviors and maximize their fitness.

## Acknowledgements

We acknowledged the helpful comments of Christian Kost, Shraddha Shitut, and Sanne Westhoff on an earlier version of this manuscript and assistance with NMR analysis from Lina Bayona and Changsheng Wu. We acknowledge Bogdan Florea of Leiden University for running and monitoring proteome measurements and the bio-organic synthesis group at Leiden University for providing access to instrumentation. Funding was provided by the China Scholarship Council (CSC) to Z.Z., by the JPI-AMR to A.L. and D.E.R. and by NWO to D.C.

## Author contributions

Z.Z., D.C. and D.R. conceptualized and planned the study. Z.Z., C. D., and F.d.B. collected data. All authors analyzed and interpreted the results. Z.Z., D.C. and D.R. drafted the manuscript with input from the other authors. All authors approved the final submitted version of this paper.

## Materials and Methods

### Bacterial strains and growth conditions

*Streptomyces coelicolor* A3(2) M145 was obtained from the John Innes centre strain collection. The strain was cultivated at 30 °C on Soy Flour Mannitol Media agar plates (SFM) for strain isolation and to quantify CFU^48^. SFM contains, per liter: 20 g mannitol, 20 g agar and 20 g soya flour. To examine antibiotic production and to extract secondary metabolites, we used minimal media (MM) supplied with 0.5% mannitol and 740 μg ml^−1^ casamino acids (CA). MM contains, per liter, 0.5 g L-asparagine, 0.5 g K_2_HPO_4_, 0.2 g MgSO_4_.7H_2_O, 0.01 g FeSO_4_.7H_2_0 and 10 g agar. For DNA extraction, strains were grown in liquid flasks shaken at 200 rpm at 30 °C in TSBS:YEME (1:1 v:v) supplemented with 0.5% glycine and 5mM MgCl_2_. TSBS contains 30 g tryptic soya broth powder and 100 g sucrose/liter. And YEME contains 3 g yeast extract, 5 g peptone, 3 g malt extract, 10 g glucose and 340 g sucrose. *E. coli* and *B. subtilis* were cultivated at 37 °C in LB media with constant shaking or on LB-agar plates.

All strains were derived from a single isolate of *S. coelicolor* A3(2) M145 (designed as WT). Briefly, samples from a frozen spore stock were diluted and plated onto SFM agar to obtain single colonies. After 5-days of growth, single colonies with WT morphology were diluted and plated onto another SFM plate. From each plate single colonies with conspicuously mutant phenotypes were picked into sterile water and plated at appropriate dilutions onto SFM agar (n = 3/colony), from which we estimated the frequency of different mutant phenotype classes. Each derived type was plated to confluence on SFM agar, and after 7 days of growth, spores were harvested to generate spore stocks which were stored at −80 °C in 20% glycerol. To quantify mutation frequency, single colonies were grown for 5 days on three different media, then we picked the colonies with WT morphology, diluted and plated them onto the corresponding media. Mutatation frequency was scored based on the phenotypes after 3 to 5 days.

### Phenotypic scoring

Two phenotypes that are related to the loss of loci in the right arm were scored (n = 3/strain). The arginine auxotrophs were identified by replicating 10^3^ CFU of each strain on MM supplied with 0.5% Mannitol with and without 37 μg ml^−1^ arginine^45^. After 5 days of growth, auxotrophs were identified as those strains that only grow on the media supplied with arginine. Chloramphenicol resistance was estimated by using the disk diffusion method. In detail, 2×10^5^ spores were spread onto MM supplemented with 740 μg ml^−1^ casamino acids ^45^ in 12 mm square petri dishes, followed by placing a paper disk containing 25 μg chloramphenicol on it. After 4 days, the radius of the inhibition zone around the disk was measured using ImageJ ^49^. Inhibition zones that were smaller than 5 mm were scored as resistant while those that are larger than 5 mm were scored as susceptible.

### Antibiotic production

Spores of each strain were diluted to 10^5^ CFU ml^−1^ in MiliQ water and 1 μl was spotted onto MM + casamino acids agar plates (n = 4/strain). After growth for 5 days at 30 °C, plates were covered with 15 ml of LB soft agar (0.7 %) containing 300 μl of a freshly grown indicator strain (OD_600_ 0.4-0.6). After overnight incubation at 30 °C, zones of inhibition around producing colonies were measured using ImageJ.

The bioactivity against *Streptomyces* isolates was tested for four strains, 2H1A, 8H1B, 9H1B and 9H1A. Three ml of SFM agar was poured onto each well of a 100mm square petri dish (Thermo Scientific, USA), after which we spotted 1 μl of each test strain containing ∼10^6^ total spores in the corner of each well. After 5-day growth, 500 μl of MM supplemented with 0.5% mannitol and 740 μg ml^−1^ casamino acids containing ∼10^5^ spores of the target strain was overlaid on top. Zones of inhibition were measured 2 days later using ImageJ.

### ^1^H NMR profiling and data analysis

2×10^5^ spores were spread onto MM + casamino acids in 12 mm square petri dishes (n=3/strain, except n = 2 for one WT clone). After 5-days of incubation at 30 °C, agar was chopped into small pieces using a sterile metal spatula and secreted compounds were extracted in 50 ml ethyl acetate for 72h at room temperature. Next, the supernatant was poured off and evaporated at 37 °C using a rotating evaporator. Pellets were obtained by drying at room temperature to remove extra solvents and then freeze-dried to remove remaining water. After adding 500 μl methanol-d_4_ to the dried pellets, the mixtures were vortexed for 30 seconds followed by a 10-minute centrifugation at 16000 rpm. The supernatants were then loaded into a 3 mm-NMR tube and analyzed using 600 MHz ^1^H NMR (Bruker, Karlsruhe, Germany)^36, 37^.

Data bucketing of NMR profiles was performed using AMIX software (version 3.9.12, Bruker Biospin GmbH) set to include the region from δ 10.02 to 0.2 with a bin of 0.04 ppm scaled to total intensity, while the signal regions of residual HDO in methanol (δ 4.9 – 4.7) and methanol (δ 3.34 – 3.28) were excluded. Multivariate data analysis was performed with the SIMCA software (version 15, Umetrics, Sweden)^36^.

### Mass spectrometry-based quantitative proteomics

10^6^ spores were spotted on SFM agar covered with cellophane and incubated for 5 days at 30 °C. Colonies were scraped off and snap frozen in liquid N_2_ in tubes, then lysed 3 times in a pre-cooled TissueLyser (QIAGEN). Proteins were dissolved in lysis buffer (4% SDS, 100 mM Tris-HCl pH7.6, 50 mM EDTA) and then precipitated using chloroform-methanol^50^. The dried proteins were dissolved in 0.1% RapiGest SF surfactant (Waters) at 95°C. Protein digestion steps were done according to van Rooden et al.^51^. After digestion, trifluoroacetic acid was added for complete degradation and removal of RapiGest SF. Peptide solution containing 8 μg peptide was then cleaned and desalted using the STAGE-Tipping technique^52^. Final peptide concentration was adjusted to 40 ng/μl with 3% acetonitrile, 0.5% formic acid solution. 200 ng of digested peptide was injected and analysed by reverse-phase liquid chromatography on a nanoAcquity UPLC system (Waters) equipped with HSS-T3 C18 1.8 μm, 75 μm X 250 mm column (Water). A gradient from 1% to 40% acetonitrile in 110 min was applied, [Glu^1^]-fibrinopeptide B was used as lock mass compound and sampled every 30 s. Online MS/MS analysis was done using Synapt G2-Si HDMS mass spectrometer (Waters) with an UDMS^E^ method set up as described^51^.

Mass spectrum data were generated using ProteinLynx Global SERVER (PLGS, version 3.0.3), with MS^E^ processing parameters with charge 2 lock mass 785.8426. Reference protein database was downloaded from GenBank with the accession number NC_003888.3. The resulting data were imported to ISOQuant^53^ for label-free quantification. TOP3 quantification result from ISOQuant was used in later data processing steps.

In total, of the 7767 proteins from the database, 2261 proteins were identified across all samples. For each sample, on average 1435 proteins were identified. TOP3 quantification was filtered to remove identifications meeting both criteria: 1. identified in less than 70% of samples of each strain; 2. The sum of TOP3 value less than 1 × 10^5^. This led to removal of 297 protein quantification results. Proteins were considered significantly altered in expression when log_2_ fold change ≧ 1 and p-value ≦ 0.05. Volcano plots were made from filtered data, with the four biosynthetic gene clusters color coded.

### CFU production

To quantify CFU, 10^4^ spores of each strain were spread onto SFM agar (n = 3/strain, except n = 2 for 9H1B) and left to grow for 7 days to confluence. After 7 days, spores were harvested by first adding 10 ml MilliQ water onto the plate and then using a cotton swab to remove spores and mycelial fragments from the plate surface. Next, the water suspension was filtered through an 18 gauge syringe plugged with cotton wool to remove mycelial fragments. After centrifuging the filtered suspension at 4000 rpm for 10 min, the supernatant was poured off and the spore pellet was dissolved in a total volume of 1ml 20% glycerol. CFU per plate was determined via serial dilution onto SFM agar.

### DNA extraction and sequencing

Nine strains, including one wild-type and eight mutants, were selected for sequencing with the Sequel system from Pacific Bioscience (PacBio, USA). Roughly 10^8^ spores were inoculated in 25 ml TSBS:YEME (1:1 v:v) supplemented with 0.5% glycine and 5 mM MgCl_2_ and cultivated at 30°C with 200 rpm shaking speed overnight. The pellet was then collected after centrifugation and washed twice with 10.3% sucrose. Samples were then resuspended in DNA/RNA shield (Zymo Research, USA) with 10x volume at room temperature and sent to be commercially sequenced at Baseclear (Leiden, The Netherlands).

Subreads of the sequenced results shorter than 50 bp were filtered and stored in BAM format. The reference alignments were performed against *Streptomyces coelicolor* A3(2) genome (NC_003888.3) using BLASR (v5.3.2) ^54^. Resulting BAM files were then sorted and indexed using SAMtools (v1.9) ^55^. For the calculation of genome rearrangements, the depths were called and exported through the depth function in SAMtools. The edges of genome were identified by manually checking the break point where the coverage drops to zero. The size of the amplified region was defined by the dramatically higher coverage compared to the adjacent sequences. All results were further confirmed by visualizing them in IGV (v2.4.15) ^56, 57^.

### Pulsed-field gel electrophoresis (PFGE)

Approximately 10^8^ spores were inoculated into 25 ml TSBS:YEME (1:1 v:v) supplemented with 0.5% glycine and 5 mM MgCl_2_ and cultivated overnight at 30 °C at 200 rpm. After centrifuging the culture at 4000 rpm for 10 min, the pellet was resuspended in 400 μl cell suspension buffer containing 100 mM Tris: 100mM EDTA at pH 8.0 and 1 mg ml^−1^ lysozyme and mixed with the same volume of 1% SeaKem Gold Agarose (Lonza, USA) in TE buffer containing 10 mM Tris: 1 mM EDTA at pH 8.0 with 1% SDS. This mixture was immediately loaded into the PFGE plug mold (Bio-rad, USA). Next, plugs were lysed in 5 ml cell lysis buffer containing 50 mM Tris: 50 mM EDTA at pH 8.0, 1% N-Lauroylsarcosine, sodium salt and 4 mg ml^−1^ lysozyme incubated for 4 hours at 37 °C with gentle agitation. This was then followed by a 5-hour incubation in 5 ml cell lysis buffer containing 0.1 mg ml^−1^ proteinase K at 56 °C and 50 rpm. Finally, the plug was washed twice in pre-heated milliQ water and 4 times in pre-heated TE buffer and incubated at 56 °C for at least 15 min with gentle mixing. Plugs were sliced into 2 mm width pieces and pre-soaked in 200 μl 1x NEB 3.1 buffer for at least 30min. After replacing the buffer with 200 μl 1x NEB 3.1 buffer, 2 μl of the rare-cutter AseI (New England Biolabs, UK) was added and incubated at 30 °C overnight. 1% agarose was used for running fragments in 0.5x freshly prepared TBE. 0.225-2.2 Mb *S. cerevisiae* chromosomal DNA (Bio-rad, USA) and wild-type *S. coelicolor* DNA were used as size markers for estimate fragment sizes. Two electrophoresis conditions were applied to separate and visualize the smaller (<1016 kb) and larger fragments (>1016 kb): (Switch time: 2.2-75 s; Voltage: 200 V; Running time: 19 h) and (Switch time: 60-125 s; Voltage: 200 V; Running time: 20 h), respectively.

PFGE results were compared to the AseI restriction maps of the wild-type strain which contains 17 fragments ranging from 26bp to 1601 kb (Figure S3). Two fragments, 240 kb and 632 kb, can be easily resolved if they are deleted from the left arm while one large 1601 kb fragment can be affected when deletions occur in right arm.

### Fitness estimates

The relative fitness of four fully sequenced mutant isolates, 2H1A, 8H1B, 9H1B and 9H1A, was estimated during pairwise competition with the wild-type parent. To distinguish strains, we first transformed mutant and wild-type strains with the integrating plasmids PIJ82 and pSET152 which confer hygromycin B and apramycin resistance, respectively. Potential fitness effects of the markers were determined by generating two WT variants that were transformed with either single marker. No effects were observed in these control experiments (One sample t-test, t = 2.029, df = 7, p = 0.082). Fitness assays were initiated by normalizing each strain to a density of 10^6^ spores/ml and then mixing at different starting ratios of Mutant:WT. 100 μl of this mixture, containing 10^5^ spores, was plated as a lawn onto SFM agar and incubated at 30 °C for 5 days, while another fraction of the sample was plated after serial dilution onto SFM containing either 50 μg ml^−1^ apramycin or 50 μg ml^−1^ hygromycin B to precisely quantify the densities of each strain. After 5 days of growth, bacteria were harvested as above and plated by serial dilution onto SFM containing either 50 μg ml^−1^ apramycin or 50 μg ml^−1^ hygromycin B. Fitness was quantified, following Lenski et al ^58^, by calculating the ratio of the Malthusian parameters of both strains. Values below 1 indicate that mutant strains have lower fitness than the WT strain. More detailed assays were carried out with strain 9H1A, where we simultaneously estimated the fitness of this strain at a broader range of frequencies from 10-99% and determined how the frequency of the mutant strain influenced antibiotic production, as measured by the size of the zone of inhibition against a *B. subtilis* indicator in a soft agar overlay.

## Supplemental Figures: Figure S1-S7

**Figure S1:**
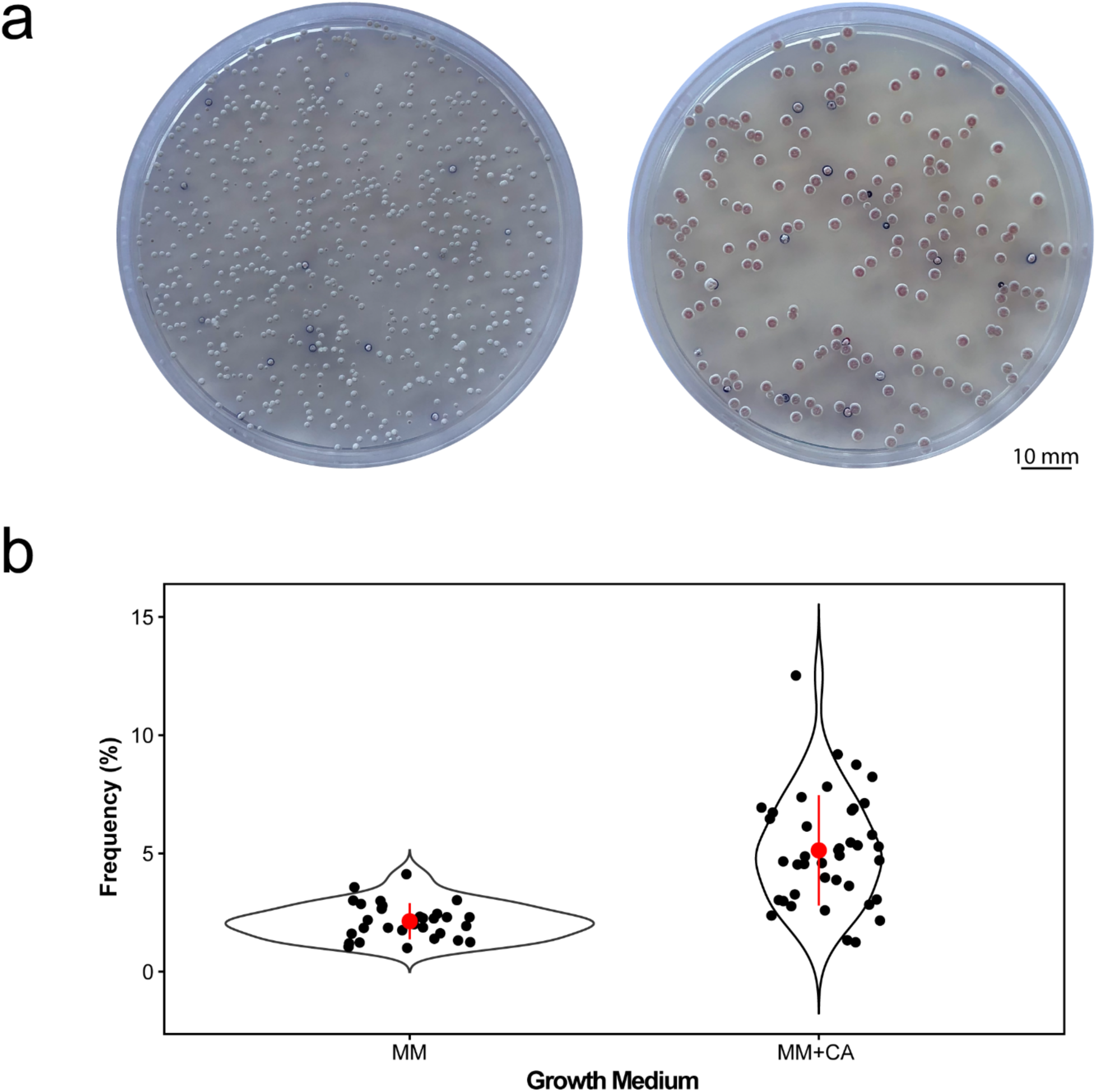
(a) Examples of mutant and wild-type colonies growing on minimal media (MM) (Left) or minimal media supplemented with casamino acids (MM+CA) (Right). (b) The frequency of mutants emerging from WT colonies on both media types.

**Figure S2:**
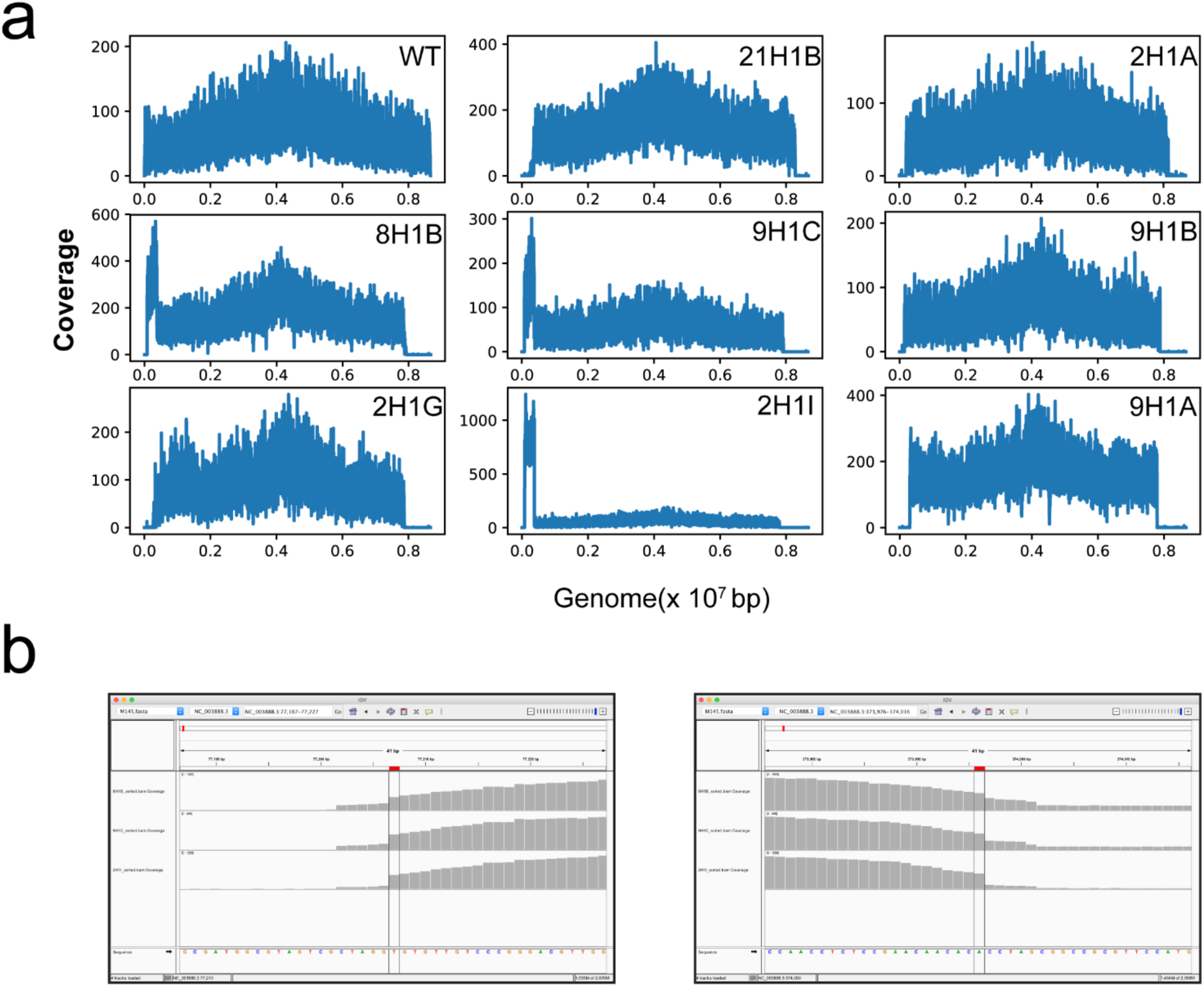
(a) PacBio reads mapped to the *S. coelicolor* M145 reference genome indicating coverage and highlighting breakpoints for genome deletions, where coverage declines to 0, as well as the size of the conserved amplified regions on the left chromosomal arms of strains 8H1B, 9H1C and 2H1I. (b) IGV snapshots of the left and right borders of amplified regions of strains 8H1B, 9H1C and 2H1I.

**Figure S3:**
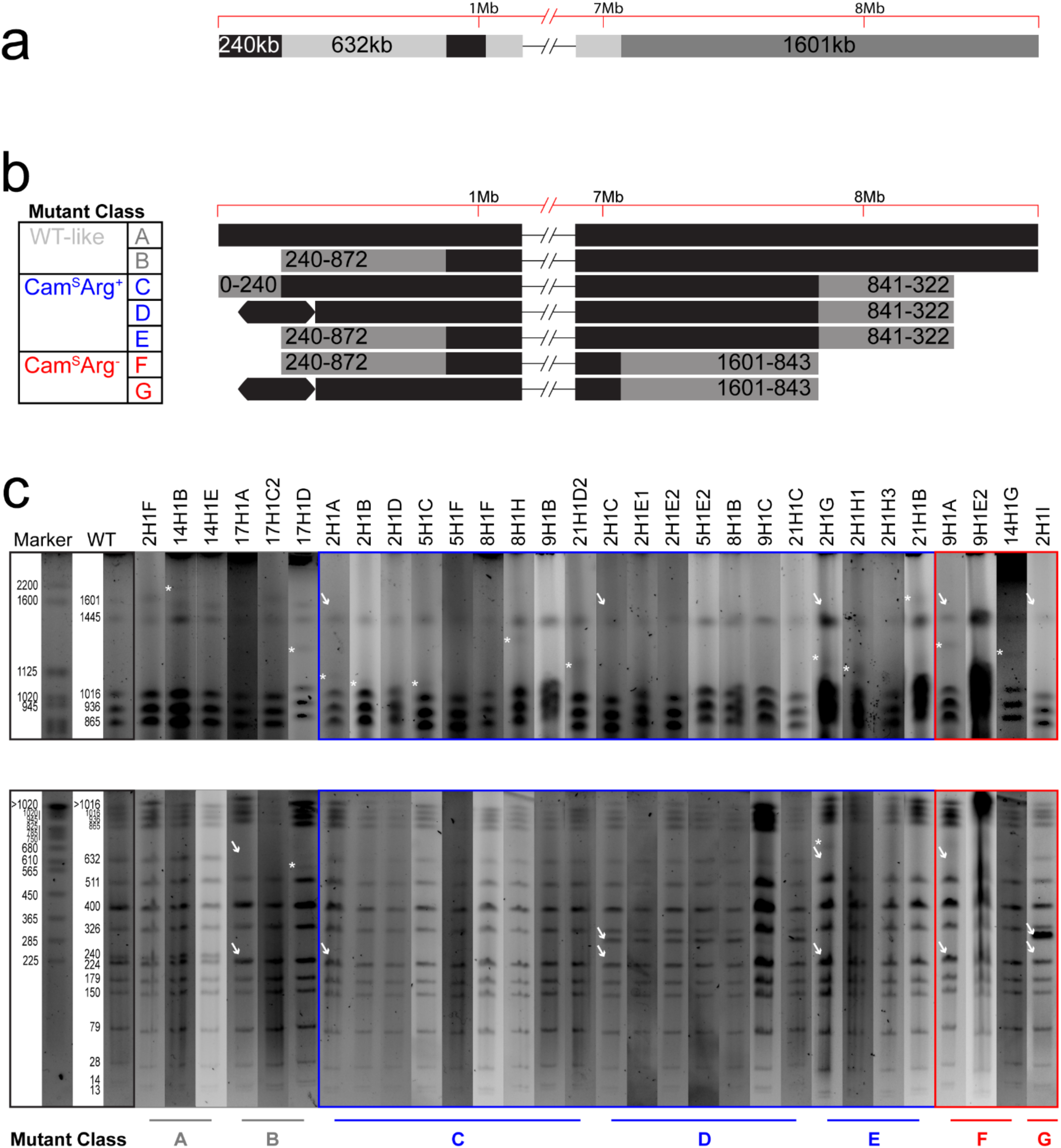
Pulse field gel electrophoresis (PFGE) results of all sampled strains. (a) The schematic of changes to *S. coelicolor* fragments in mutant isolates after AseI digestion. Two bands (240 kb and 632 kb) and one 1601 kb band can be affected on the left and right arms, respectively. (b) Schematic adjusted from Figure 2b with more detailed mutant classes, designated A-G. (c) PFGE results of 30 sampled strains. Two running conditions are used to visualize larger (top panel) or smaller (bottom panel) fragments. (Detailed running conditions are given in the Materials and Methods). White arrows indicate missing or newly appearing of the bands for different mutant class. Asterisks indicate the new bands that were used to evaluate precise genome length.

**Figure S4:**
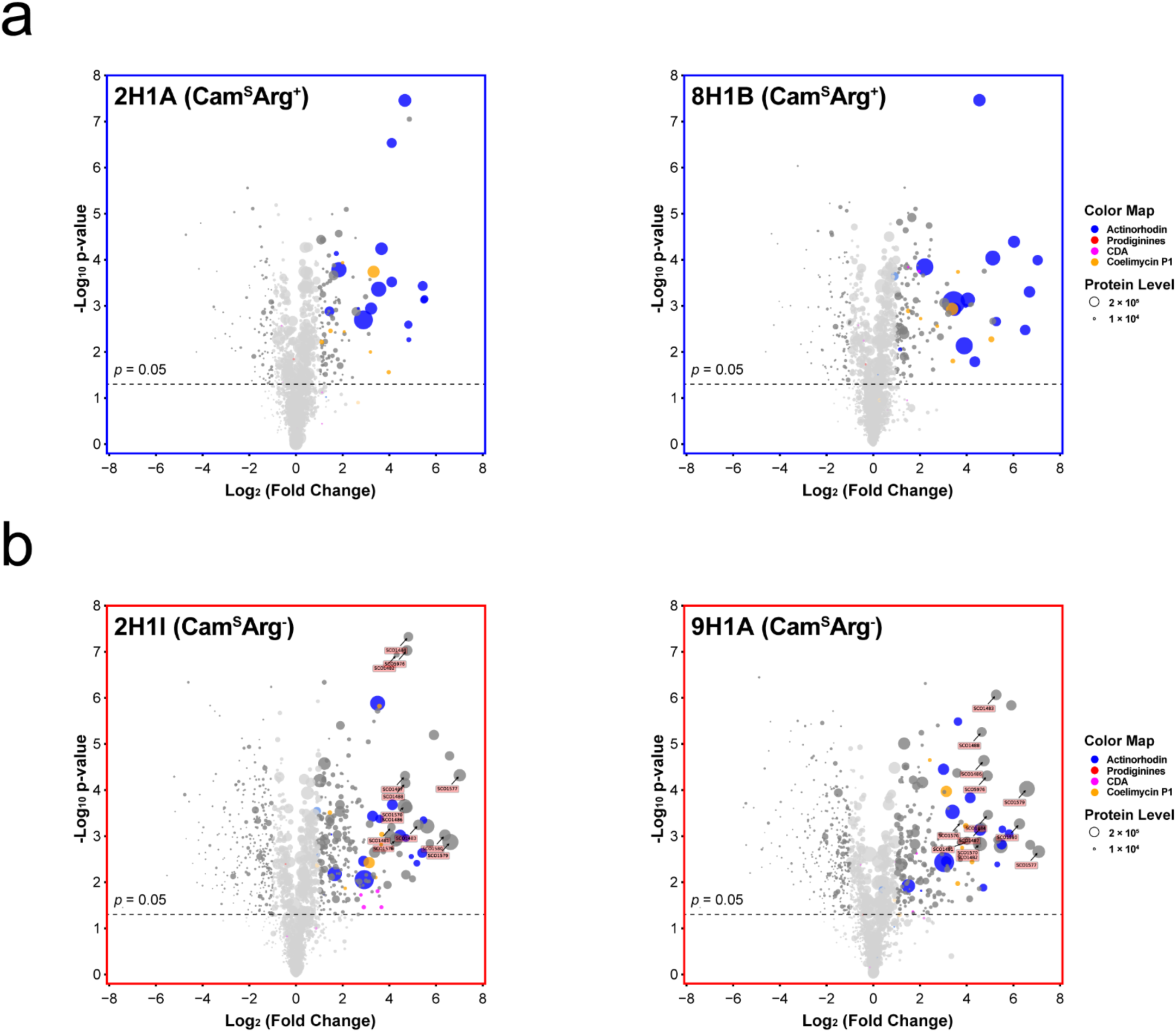
Volcano plots of proteomics from two Cam^S^Arg^+^ strains (a) and two Cam^S^Arg^−^ strains with annotated genes from arginine and pyrimidine biosynthesis pathways (b).

**Figure S5:**
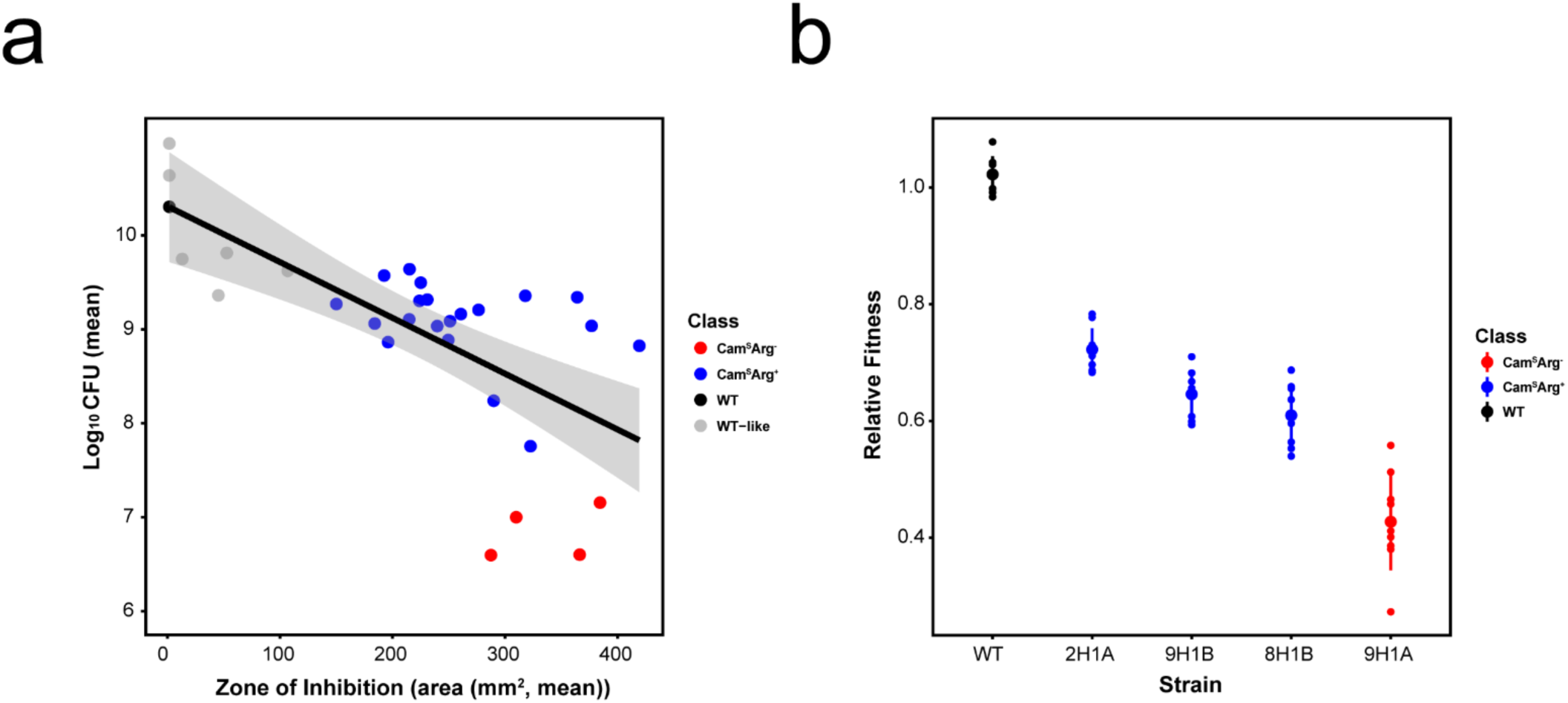
(a) The trade-off between antibiotic production and reproductive capacity, partitioned by different mutant classes. (b) Relative fitness of four selected strains. Detailed methods are given in the Methods and Materials.

**Figure S6:**
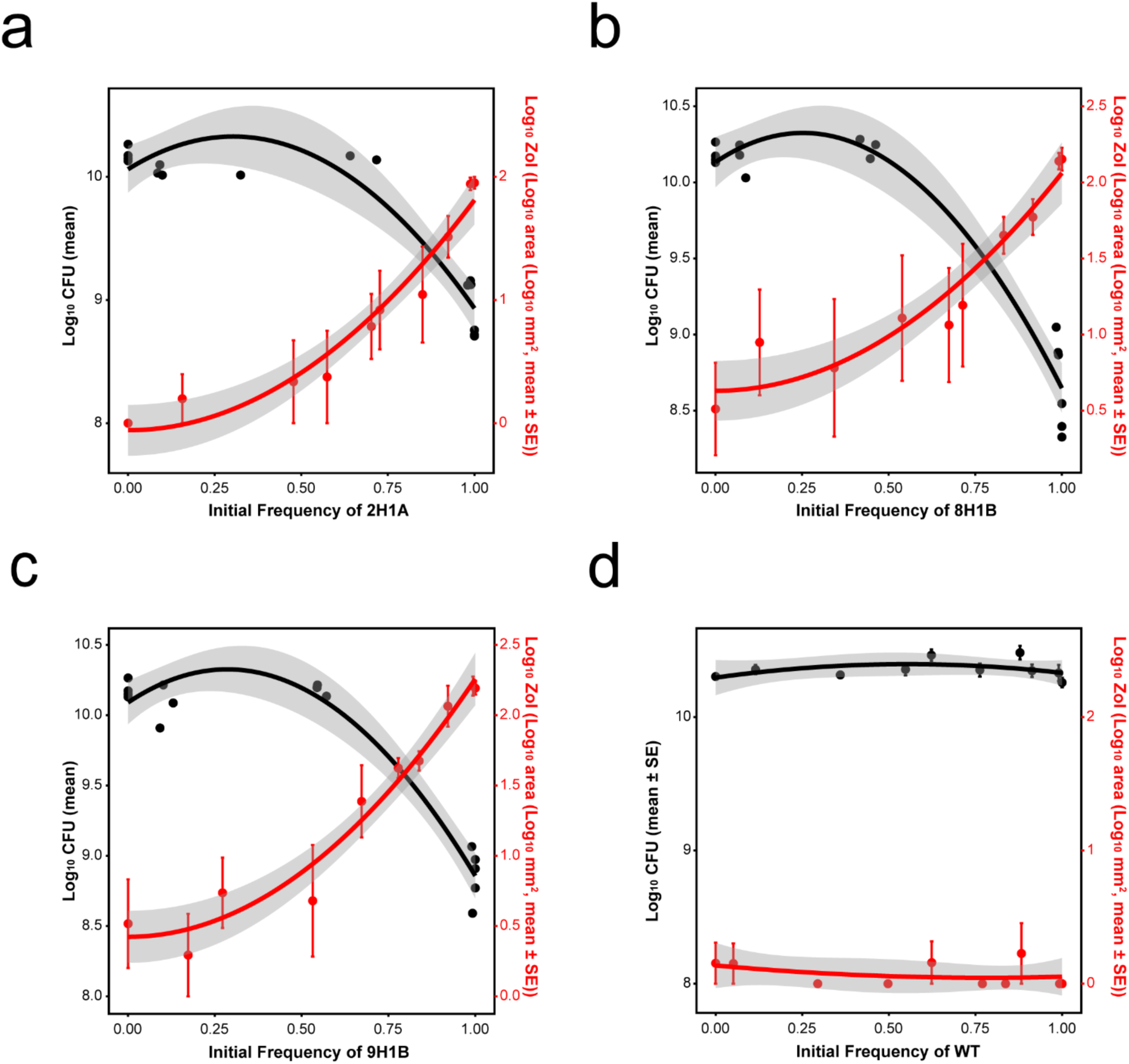
Extended evidence for division of labour during co-culture of the WT and three mutant strains at different starting frequencies. Increasing frequencies of mutants cause increased antibiotic production (red) for (a) 2H1A (F_1,8_ = 170.3, r^2^ = 0.955, *p* < 0.001), (b) 8H1B (F_1,8_ = 105.3, r^2^ = 0.929, *p* < 0.001) and (c) 9H1B (F_1,8_ = 201.1, r^2^ = 0.962, *p* < 0.001) but not (d) WT (F_2,7_ = 0.576, r^2^ = 0.141, *p* = 0.587). Increasing frequencies of mutants only negatively impact colony fitness at frequencies > ∼ 50% (black) for (a) 2H1A (F_2,13_ = 59.44, r^2^ = 0.901, *p* < 0.001), (b) 8H1B (F_2,13_ = 131.7, r^2^ = 0.953, *p* < 0.001) and (c) 9H1B (F_2,12_ = 201.1, r^2^ = 0.944, *p* < 0.001) but not (d) WT (F_2,7_ = 1.076, r^2^ = 0.235, *p* = 0.391). Quadratic regression lines include the 95% CI.

**Figure S7:**
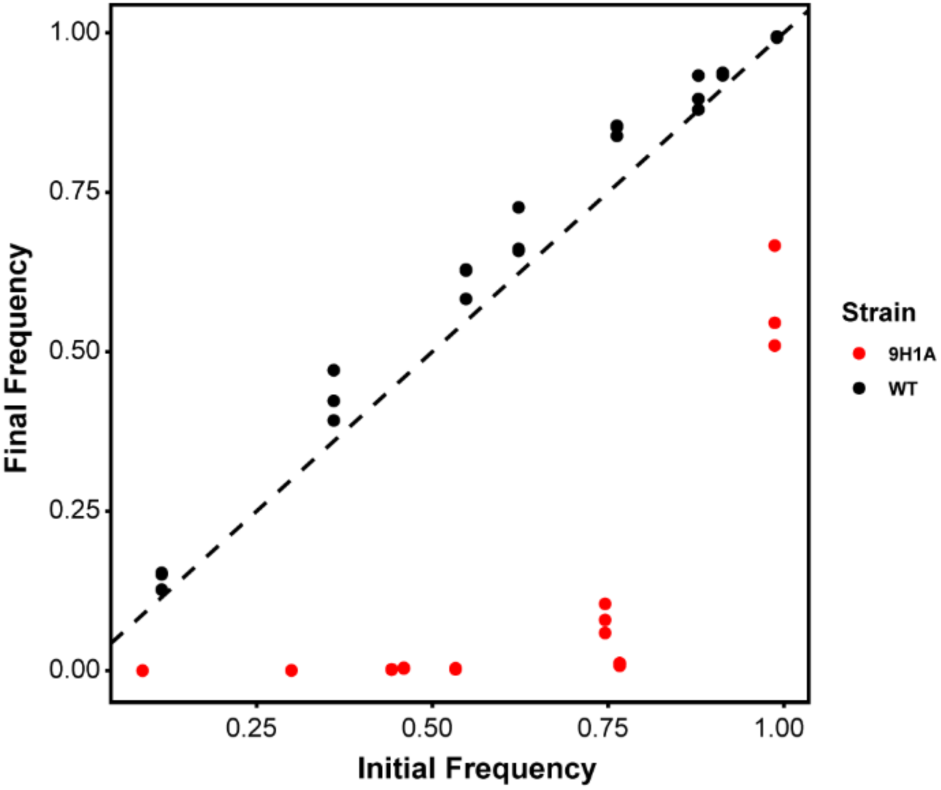
Initial and final frequencies of strains during pairwise competition assays between 9H1A (red) and the wild-type or between two differentially marked wild-type strains (black). The dashed line indicates that initial and final frequencies are equal, while values below the line indicate that the competing strain has declined during the competition assay. While the differentially marked wild-type strains have equal fitness, the mutant strain has dramatically reduced fitness at every starting frequency.

